# Novel Dopamine 4 Receptor Ligands Differentially Ameliorate ADHD-like Behaviors in Spontaneously Hypertensive Rats

**DOI:** 10.1101/2025.11.26.690786

**Authors:** Ike C. de la Peña, Samantha Andino, Ashley Amis, Mohammad A. Alkhatib, Tian Li, Thomas M. Keck, Comfort A. Boateng

**Author notes:** Corresponding Authors: Ike C. de la Peña- Department of Pharmaceutical and Administrative Sciences, Loma Linda University School of Pharmacy, Loma Linda, California, 92350, United States, Thomas M. Keck − Department of Chemistry and Biochemistry, Department of Biological and Biomedical Sciences, College of Science and Mathematics, Rowan University, Glassboro, New Jersey 08028, United States, Comfort A. Boateng − Department of Basic Pharmaceutical Sciences, Fred Wilson School of Pharmacy, High Point University, High Point, North Carolina 27268, United States.

## Abstract

**Rationale:** Dopamine D4 receptors (D4Rs) have been implicated in the pathophysiology of attention-deficit/hyperactivity disorder (ADHD), yet their precise role and therapeutic relevance remain underexplored. Highly selective D4R compounds may provide a valuable tool to elucidate D4R function and assess their potential as non-stimulant ADHD treatments.

**Objectives:** This study examined the behavioral effects of two novel D4R drugs, namely, FMJ-01-38 (high-efficacy partial agonist) and FMJ-01-54 (full antagonist) in adolescent spontaneously hypertensive (SHR/NCrl) rats, a validated ADHD model, and Wistar controls.

**Methods:** Rats received intraperitoneal FMJ-01-38 or FMJ-01-54 (5–10 mg/kg) or vehicle prior to behavioral assays assessing locomotor activity (open field tests), recognition memory (novel object preference), attention and working memory (Y-maze test), and impulsivity (delay discounting task).

**Results:** FMJ-01-38 dose-dependently reduced locomotor hyperactivity and improved spontaneous alternation behavior in SHR/NCrl; at 5 mg/kg it enhanced novel-object preference and decreased impulsive choice and action, indicating attenuation of ADHD-like symptoms and cognitive enhancement. FMJ-01-54 produced similar improvements in Y-maze and novel-object performance without altering locomotor activity or impulsivity of SHR/NCrl, suggesting selective cognitive improvement. In Wistar rats, FMJ-01-38 increased novel-object preference only at the 5 mg/kg dose, while FMJ-01-54 treatment did not produce any significant behavioral effects.

**Conclusions:** These findings demonstrate that D4R modulation, through either partial agonism or antagonism, differentially ameliorates ADHD-related behaviors. Both FMJ-01-38 and FMJ-01-54 produced minimal effects in control animals, suggesting pathology-specific efficacy and highlighting D4R ligands as promising non-stimulant therapeutic candidates for ADHD.

## Introduction

Attention-deficit/hyperactivity disorder (ADHD) is a prevalent neurodevelopmental, childhood-onset psychiatric disorder characterized by age-inappropriate inattention, impulsiveness, and hyperactivity, with significant social, academic, and occupational difficulties continuing to adulthood (Faraone et al., 2024; Thapar and Cooper, 2016). Pharmacotherapy for ADHD has long relied on psychostimulant medications such as methylphenidate and amphetamine (Faraone, 2018), drugs with potential for abuse, misuse, and diversion (Forrest et al., 2025; Wilens et al., 2008). Additionally, up to one-third of patients exhibit a suboptimal response to stimulant therapy (Childress and Sallee, 2014), highlighting the need for alternative, preferably non-stimulant treatment options for ADHD.

Several preclinical and clinical studies suggest the involvement of the dopamine D4 receptor (D4R) in ADHD (Ferré et al., 2022; González et al., 2012; Oak et al., 2000; Ptácek et al., 2011; Qin et al., 2016) and in the response to psychostimulant drugs used to treat the disorder (Cheon et al., 2007; Keck et al., 2013; Michaelides et al., 2010; Naumova et al., 2019). D4Rs have high affinity for dopamine and are abundantly expressed in fronto-striatal pathways linking the striatum and prefrontal cortex (PFC) (Meador-Woodruff et al., 1996; Tomlinson et al., 2015), brain regions that regulate critical processes such as attention, executive function, and behavioral control. Alterations in PFC activity and cortico-striatal circuitry have been well documented in ADHD (Arnsten, 2009; Brennan and Arnsten, 2008; Aki Nikolaidis et al., 2022). Evidence indicates that deficient D4R function is strongly linked to hyperactivity and inattention (Erlij et al., 2012; Kegel and Bus, 2013; Lasky-Su et al., 2008; Tovo-Rodrigues et al., 2014), while D4R dysfunction correlated with poor performance on measures of intelligence and working memory in individuals with ADHD (Altink et al., 2012; Kebir et al., 2009; Loo et al., 2008).

Pharmacological targeting of D4R has yielded promising results regarding the potential of D4R drugs as ADHD treatments. For instance, selective D4R antagonists such as L-745,870 and U-101,958 reduced hyperactivity in dopamine-depleted and lesion-based ADHD animal models while not affecting locomotor activity in control rats (Zhang et al., 2002). D4R agonists, including ABT-724, improved spatial learning and reduced hyperactivity in the spontaneously hypertensive rats (SHR) ADHD model (Yin et al., 2014), whereas A-412997 enhanced recognition memory and cognitive performance in both SHR and unimpaired, i.e., non-ADHD rats (Browman et al., 2005; Kocsis et al., 2014; Woolley et al., 2008).

Nevertheless, some D4R agonists have also been reported to produce undesirable behavioral and physiological effects, including locomotor activation (Nayak and Cassaday, 2003; Woolley et al., 2008), exacerbation of amphetamine-induced hyperactivity or psychosis-like behavior (Chestnykh et al., 2023), and induction of penile erection (Brioni et al., 2004; Patel et al., 2006). Moreover, a previous clinical trial using the selective D4R antagonist L-745,870 (also known as MK-0929) in adults with ADHD did not show a significant effect compared to placebo (Rivkin et al., 2011). These results highlight the need for further research and drug development to optimize the therapeutic potential of D4R modulation for ADHD. Partial agonists, such as RO-10-5824, enhanced exploratory behaviors in mice without affecting locomotion (Powell et al., 2003) and increased PFC gamma band activity common marmosets, suggesting improved attention and impulse-control (Nakazawa et al., 2015). However, the efficacy of this compound, and more broadly the role of D4R partial agonists in ADHD animal models, has not yet been evaluated.

In previous studies, we reported the synthesis and pharmacological characterizations of several novel D4R ligands based on the classical D4R partial agonist A-412997 (Boateng et al., 2023; Keck et al., 2019). Compounds were profiled using radioligand binding displacement assays, β-arrestin recruitment assays, cyclic AMP inhibition assays, and molecular dynamics computational modeling. We identified several novel D4R-selective compounds (K_i_ ≤ 4.3 nM and >100-fold binding selectivity vs. other D2-like receptors) with diverse partial agonist and antagonist profiles. Recently, we developed new analogs with improved pharmacokinetics suitable for rat behavioral studies (Alkhatib et al., 2025). In this study, we tested the effectiveness of a high-efficacy partial agonist (FMJ-01-38) and an antagonist (FMJ-01-54) in ameliorating ADHD-like behaviors in adolescent SHR/NCrl. As validated models of ADHD, SHR/NCrl display hallmark behavioral phenotypes including hyperactivity, sustained attention deficits and impulsivity relative to control strains such as the Wistar or Wistar-Kyoto (WKY) rat (dela Peña et al., 2015; Kim et al., 2024; Sagvolden et al., 2009). Moreover, SHRs showed impairment in short-term associative memory in the novel object recognition test compared to Wistar rats (Tchekalarova et al., 2023). SHR/NCrl also exhibit significantly lower *Drd4* mRNA expression and reduced D4R protein synthesis in the PFC (Li et al., 2007), aligning with the cortical dopamine dysregulation proposed in human ADHD. Our study aims to contribute to a broader understanding of the role of D4Rs in ADHD pathophysiology, and to assess the therapeutic potential of novel D4R ligands as non-stimulant ADHD treatments.

## Materials and methods

### Animals

Adolescent male SHR/NCrl and Wistar rats (4 weeks old at study onset) were obtained from Charles River Laboratories (Wilmington, MA, USA) and pair-housed (except during delay discounting testing) under controlled environmental conditions (12:12 h light/dark cycle; lights on at 6:00 a.m.; temperature 22 ± 2 °C) with *ad libitum* access to food and water except during behavioral experiments. Rats were allowed to acclimate to the housing environment for one week before testing. All procedures were conducted in accordance with the Public Health Service Policy on Humane Care and Use of Laboratory Animals and the Guide for the Care and Use of Laboratory Animals and were approved by the Loma Linda University Institutional Animal Care and Use Committee (IACUC #24-015).

### Drugs

The synthesis and characterization of the FMJ compounds, including their *in vitro* and *in vivo* pharmacokinetic and pharmacodynamic properties, have been described previously (Keck et al., 2019; Alkhatib et al., 2025). All compounds were dissolved in 20–30% DMSO and 0.9% saline to achieve the desired concentrations, as detailed in earlier reports (Keck et al., 2019; Boateng et al., 2023). In the present study, doses of 5 and 10 mg/kg were used, based on results from preliminary studies (Keck et al., 2019; Alkhatib et al., 2025).

### Behavioral studies

Two cohorts of SHR/NCrl and Wistar rats were used in this study. The first cohort (*n* = 72 rats) underwent a series of behavioral assessments, namely the open field test, novel object preference test, and Y-maze task, with 24-hour intervals between each test to minimize potential carryover effects. The second cohort (*n*= 120 rats), which was subjected to food restriction and task-specific training, was used exclusively for the delay discounting task. The effects of vehicle (control) or drugs were assessed during the behavioral tests as described below. All experiments were conducted by an investigator blinded to the treatment conditions. Between sessions, the behavioral apparatuses were thoroughly cleaned with Quatricide to eliminate residual odors and prevent cross-contamination.

### Open field test: locomotor activity

The open field test is a well-established method for assessing spontaneous locomotor behavior in rodents. The procedures of the open field tests followed previously reported protocols (de la Peña et al., 2021) with some modifications. Briefly, after intraperitoneal (i.p.) injection with either vehicle, FMJ-01-38, or FMJ-01-54, each rat was placed individually and immediately in an activity chamber (47 × 47 × 50 cm), and locomotion was recorded for 30 minutes using an automated tracking system (Ethovision System, Noldus I.R., Wageningen, The Netherlands). Prior work from our group demonstrated that locomotor activity in SHR/NCrl peaked within the first 30 minutes and subsequently declined (dela Peña et al., 2021), supporting the selection of this time window for assessing drug-induced changes in locomotor activity. Total distance traveled (in centimeters), and movement duration (in seconds) were quantified to evaluate drug effects on locomotor activity of animals.

### Novel object preference tests: attention and recognition memory

The novel object preference test was used to evaluate attention and recognition memory, behavioral domains commonly altered in ADHD animal models (Kim et al., 2024; Russell et al., 2005). Testing was conducted in an activity chamber (47 × 47 × 50 cm) equipped with an overhead camera and spanned three consecutive days: habituation, familiarization, and novel object testing (de la Peña et al., 2025). On Day 1, rats were allowed to freely explore the empty arena for 10 minutes. On Day 2, two identical red cone-shaped objects were placed in the arena, and rats were allowed to explore for 10 minutes. On Day 3, one familiar object was replaced with a novel object differing in both shape and color. Drug or vehicle treatments were administered 20 minutes before the novel object testing session, which lasted 5 minutes. Exploration time, defined as sniffing or touching the object within 2 cm, was recorded manually by an experimenter blinded to treatment conditions. The percent preference for the novel object was calculated as [time exploring novel / (time exploring novel + time exploring familiar)] × 100. A reduced preference for the novel object was interpreted as impaired attention or recognition memory.

### Y-maze test: inattention/working memory

Spontaneous alternation behavior in the Y-maze task reflects both attentional and working memory processes (Sarter et al., 1988). Deficits in attention are manifested as a reduced percentage of spontaneous alternation (de la Peña, et al., 2015; Kishikawa et al., 2014). The Y-maze procedure followed methods described previously (de la Peña et al., 2015; de la Peña et al., 2021; de la Peña et al., 2025). Rats received intraperitoneal injections of vehicle or test drug 20 minutes before testing. Each animal was placed at the end of one arm and allowed to explore freely for 8 minutes. Arm entries were recorded manually, with an entry defined as placement of all four paws and the tail within an arm. From these data, the number of actual alternations (successive entries into all three arms in overlapping triplets) and the maximum possible alternations (total arm entries − 2) were determined (de la Peña et al., 2015; dela Peña et al., 2021; de la Peña et al., 2025). The percentage of spontaneous alternation, calculated as *(actual / maximum alternations) × 100*, served as the primary index of attention and working-memory functions.

### Delay discounting task (DDT): impulsivity

Delay discounting, defined as the preference for a small immediate reward over a larger delayed one, serves as an index of impulsivity (Odum, 2011). The DDT procedures followed our previous protocols (de la Peña et al., 2017; de la Peña et al., 2021). Testing was conducted in standard operant chambers (Coulbourn Instruments, Allentown, PA, USA) equipped with two levers, chamber and magazine lights, and a pellet dispenser. Prior to tests, rats were food restricted (to increase their motivation to work for food delivery) and trained to press a lever for a contingent sucrose pellet reward (45 mg, TestDiet, IN, USA). During 1-week training (30 min/day), pressing the “small and immediate” lever (L1) delivered one pellet, while the “large and delayed” lever (L5) delivered five pellets. The chamber light illuminated for 1 s before pellet delivery, followed by a 25-s timeout signaled by the magazine light. Once rats selected L5 on at least two-thirds of trials, the testing phase began. Testing followed the same procedure except that delays (0, 10, and 30 s; 3 consecutive days) were imposed before L5 reward delivery. During the delay periods, the chamber light remained illuminated, and any additional lever presses were recorded but not reinforced. The 30-min DDT session commenced 20 min after intraperitoneal administration of vehicle or drug. Total responses on both L5 and L1 levers were recorded for analysis. To maintain motivation for food reinforcement, rats remained on restricted diets throughout the study. From the DDT data, the percentage choice for the large (L5) reinforcer was calculated as an index of impulsive choice (Winstanley et al., 2006), while responses on the L1 lever during time-out periods, when no reward was available, served as a measure of impulsive action (Zeeb et al., 2016).

In previous DDT studies (de la Peña et al., 2017; de la Peña et al., 2021), we demonstrated consistent impulsive-like behavior in SHR/NCrl and Wistar rats, relative to the Wistar Kyoto rat (WKY/NCrl), genetic control strain for the SHR/NCrl. As these baseline strain differences have been well established, WKY/NCrl rats were not included in the present study allowing us to focus on the main objective of examining the effects of novel D4R-targetting drugs on impulsivity in SHR/NCrl and Wistar rats, while also adhering to the principle of reduction in animal research by minimizing unnecessary replication of known findings.

### Data Analyses

Data were analyzed using two-way ANOVA followed by pre-planned Dunnett’s or Šidák *post hoc* tests where appropriate. Dunnett’s test was used to compare each treatment group against the control, while Šidák’s test was applied for pairwise comparisons among multiple groups. Statistical analyses were conducted using GraphPad Prism Version 9.5 software (San Diego, CA, USA). Results from the above analyses were presented as the means ± S.E.M. A *p* value of < 0.05 was regarded as significant.

## Results

### FMJ-01-38, but not FMJ-01-54, reduced locomotor hyperactivity in SHR/NCrl

Figure 1A–B shows the effects of high efficacy D4R partial agonist FMJ-01-38 on the locomotor activity of SHR/NCrl and Wistar rats. A two-way ANOVA of the 30-min total distance moved revealed significant main effects of strain [F (1,11) = 148.20, p < 0.001], drug treatment [F (2,22) = 25.50, p < 0.001], and a strain × treatment interaction [F (2,22) = 8.37, p < 0.01] (Fig. 1A). *Post hoc* comparisons indicated that SHR/NCrl exhibited higher locomotor activity than Wistar rats (p < 0.001). Moreover, FMJ-01-38 treatment dose-dependently (5–10 mg/kg, i.p.) reduced the locomotor activity of SHR/NCrl, with no significant effects in Wistar rats (Fig. 1A). A two-way ANOVA of movement duration similarly revealed significant main effects of strain [F (1,11) = 150.0, p < 0.001], drug treatment [F (2,22) = 22.0, p < 0.001], and a strain × treatment [F (2,22) = 8.43, p < 0.01] interaction. *Post hoc* analysis showed that SHR/NCrl had longer movement durations than Wistar rats (p < 0.001), and that FMJ-01-38 significantly reduced movement duration in SHR/NCrl only at the 10 mg/kg (i.p.) dose (p < 0.001, Fig. 1B).

**Figure 1.**
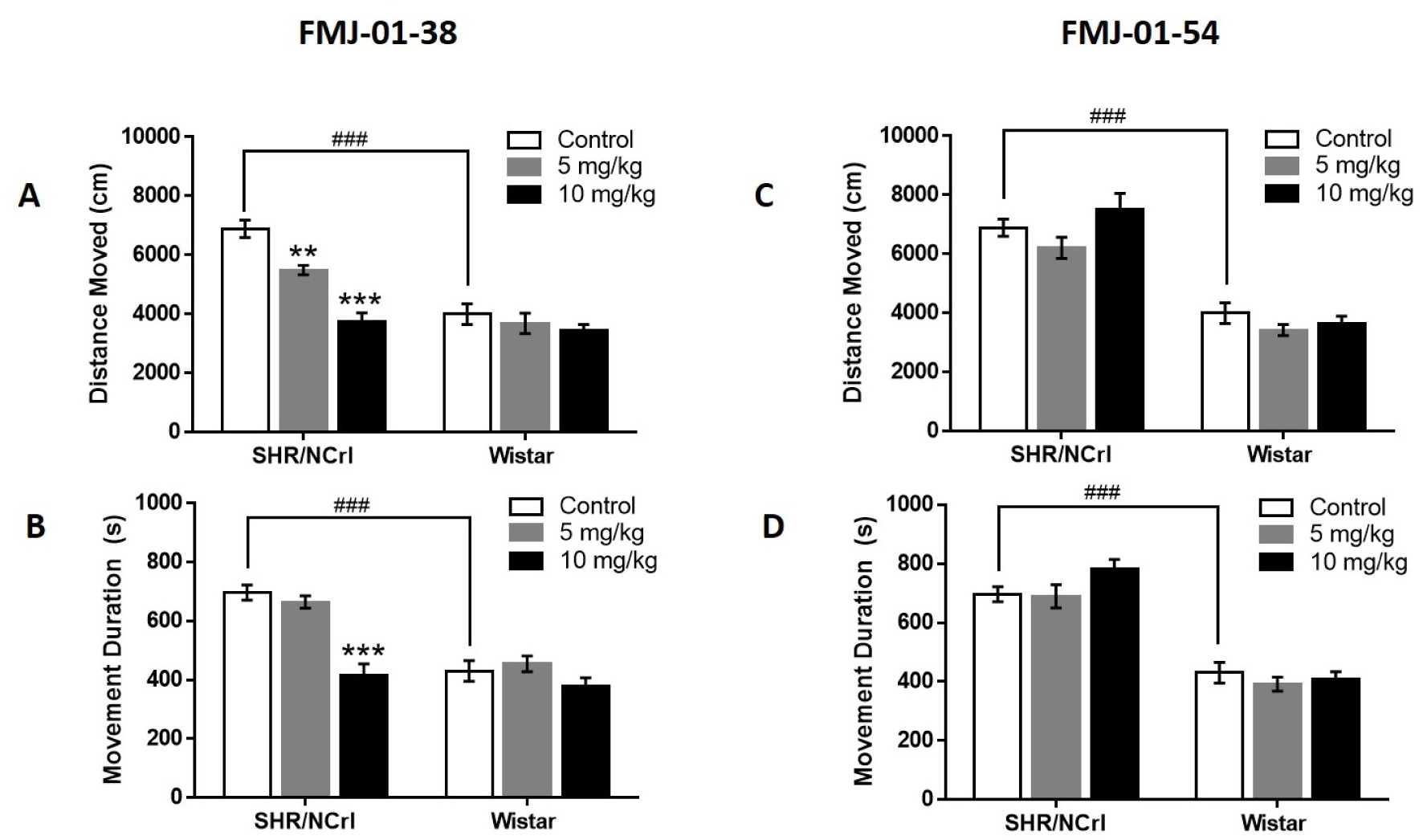
Effects of FMJ-01-38 and FMJ-01-54 on locomotor activity in adolescent SHR/NCrl and Wistar rats. Panels A and C show total distance moved (cm), and Panels B and D show movement duration (s) in the open-field test. Data are expressed as mean ± S.E.M. (n = 12 per group). ### p < 0.001; *** p < 0.001, ** p < 0.01 vs. SHR/NCrl control (vehicle-treated) rats.

Figure 1C–D depicts the effects of D4R antagonist FMJ-01-54 on locomotor activity in the same strains. Two-way ANOVA of total distance moved revealed a significant main effect of strain [F (1,11) = 76.71, p < 0.001], and drug treatment [F (2,22) = 3.55, p < 0.05) but no interaction between strain and drug treatment [F (2,22) = 1.09, p = 0.35, Fig. 1C]. Analysis of movement duration showed a significant main effect of strain [F (1,11) = 73.86, p < 0.001], but no effect of drug treatment [F (2,22) = 2.41, p = 0.11, Fig. 1D]. Consistent with these findings, SHR/NCrl had higher locomotor activity than Wistar rats (p < 0.001), whereas FMJ-01-54 produced no detectable effects on either total distance moved or movement duration in either strain across doses (Fig. 1D).

### FMJ-01-38 and FMJ-01-54 enhanced novel object preference in SHR/NCrl

Figure 2 shows the percentage of novel object preference in SHR/NCrl and Wistar rats and the effects of FMJ-01-38 and FMJ-01-54. For FMJ-01-38 (Fig. 2A), two-way ANOVA of the % novel object preference revealed significant main effects of strain [F (1, 11) = 71.74, p < 0.001] and drug treatment [F (2, 22) = 13.50, p < 0.001], with no significant strain × treatment interaction [ F (2,22) = 0.05, p = 0.95]. As expected, SHR/NCrl showed lower preference for the novel object than Wistar rats (p < 0.001), consistent with impaired recognition memory and inattention. Treatment with FMJ-01-38 (5 mg/kg, i.p.) significantly increased (p < 0.01) novel object preference in SHR/NCrl. Interestingly, the same dose also enhanced novel object preference in Wistar rats (p < 0.01), suggesting improvement of object recognition/attention in the “normal” rat strain.

**Figure 2.**
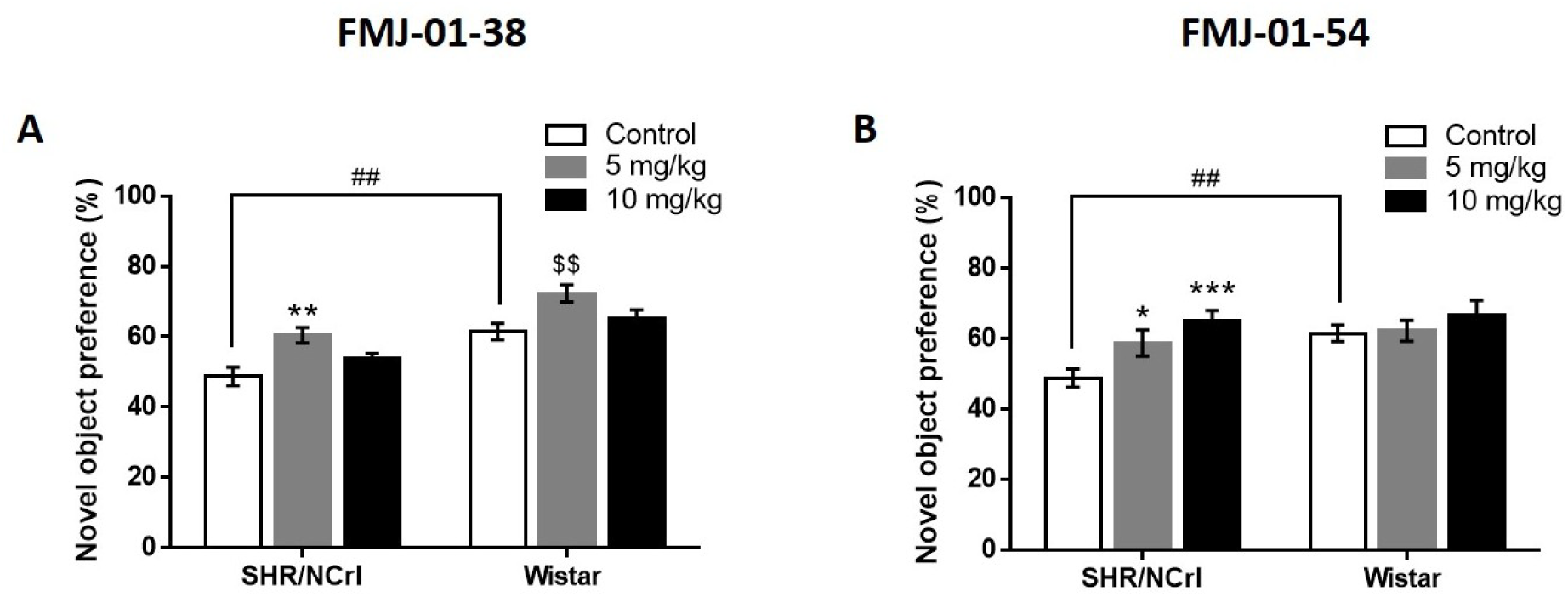
Effects of FMJ-01-38 and FMJ-01-54 on attention/recognition memory in adolescent SHR/NCrl and Wistar rats. Percent preference for the novel object (A and B) is shown. Data are expressed as mean ± S.E.M. (n = 12 per group). ## p < 0.01; ***p<0.001; ** p < 0.01, * p < 0.05 vs. SHR/NCrl control (vehicle-treated) rats; $$p<0.01 vs Wistar control (vehicle-treated) rats.

For FMJ-01-54 (Fig. 2B) two-way ANOVA likewise showed significant main effects of strain [F (1,11) = 14.15, p <0.01] and drug treatment [F (2,22) = 5.38, p < 0.05) but no significant interaction between strain and treatment [F (2,22) = 3.24, p = 0.06]. FMJ-01-54 increased the % novel object preference in SHR/NCrl in a dose-dependent manner (5–10 mg/kg, i.p.) and did not produce significant effects in Wistar rats.

### FMJ-01-38 and FMJ-01-54 improved spontaneous alternation behavior in SHR/NCrl

In line with our previous findings (de la Peña et al., 2015; de la Peña et al., 2021), SHR/NCrl showed impaired spontaneous alternation behavior compared with Wistar rats (Fig. 3A,C). A two-way ANOVA of the percentage of spontaneous alternation revealed significant main effects of strain [F (1,11) = 10.08, p < 0.01] and drug treatment [F (2,22) = 17.44, p < 0.001], but no significant strain × treatment interaction [F (2,22) = 3.02, p = 0.06]. *Post hoc* analysis showed that FMJ-01-38 (5–10 mg/kg, i.p.) dose-dependently improved spontaneous alternation behavior in SHR/NCrl (Fig. 3A). Analysis of total arm entries revealed significant main effects of strain [F (1,11) = 8.50, p < 0.05] and drug treatment [F (2,22) = 4.54, p < 0.05], with no significant strain × treatment interaction [F (2,22) = 1.39, p = 0.27] (Fig. 3B). *Post hoc* comparisons indicated that FMJ-01-38 significantly reduced total arm entries in Wistar rats at both 5 mg/kg and 10 mg/kg doses (p < 0.05).

**Figure 3.**
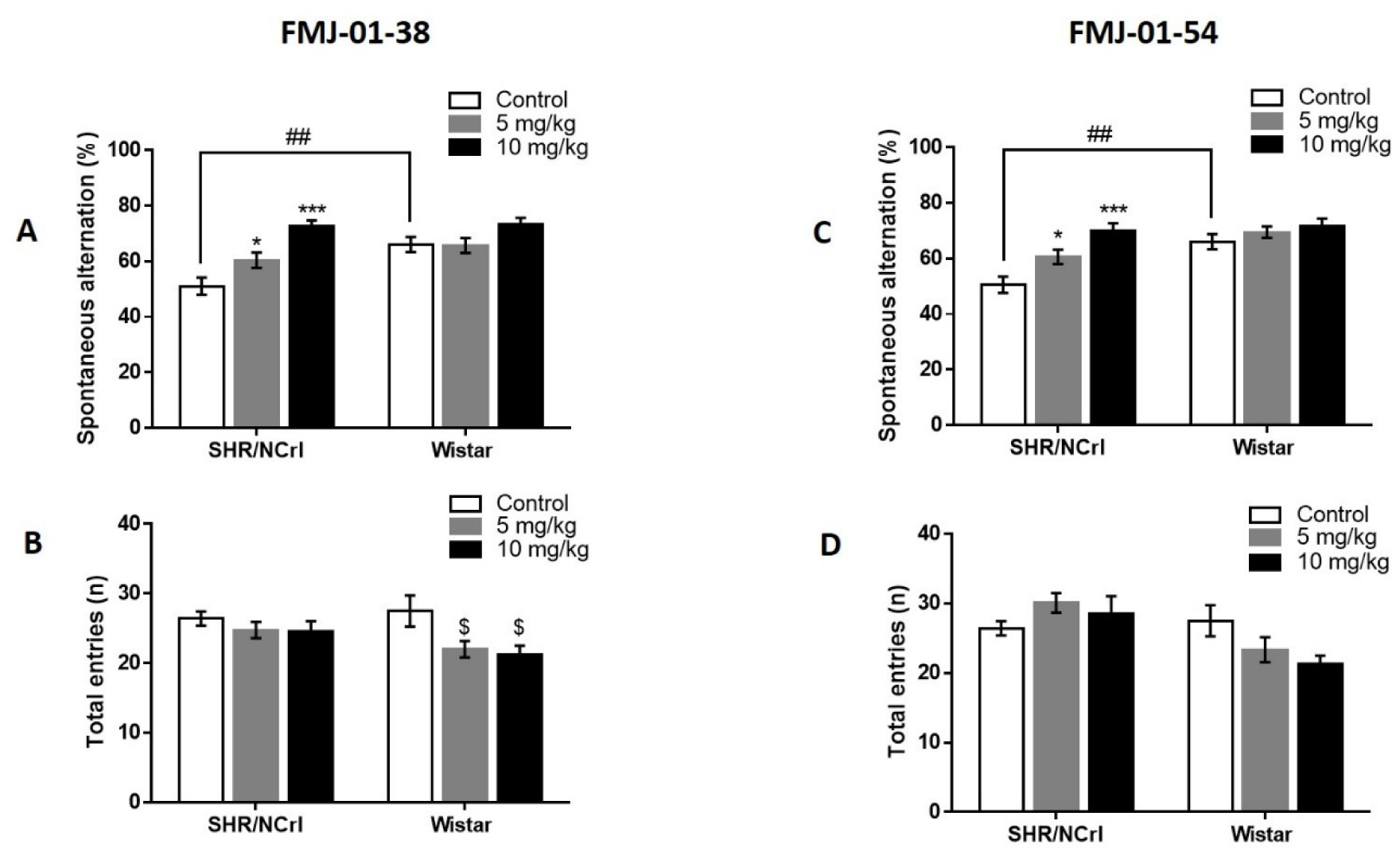
Effects of FMJ-01-38 and FMJ-01-54 on inattention-like behavior/working memory deficits in adolescent SHR/NCrl. Panels A and C show percent spontaneous alternation, and Panels B and D show total arm entries in the Y-maze test. Data are expressed as mean ± S.E.M. (n = 12 per group). ## p < 0.01; *** p < 0.001; *p < 0.05 vs. SHR/NCrl control (vehicle-treated) rats; $p < 0.05 vs. Wistar control (vehicle-treated) rats.

Regarding FMJ-01-54, two-way ANOVA of spontaneous alternation data revealed significant main effects of strain [F (1,11) = 20.93, p < 0.001] and drug treatment [F (2,22) = 13.52, p = 0.001], with no significant strain × treatment interaction [F (2,22) = 2.09, p = 0.14]. *Post hoc* analysis showed that FMJ-01-54 (5–10 mg/kg, i.p.) dose-dependently improved spontaneous alternation behavior in SHR/NCrl but not in Wistar rats (Fig. 3C). Two-way ANOVA of total arm entries for FMJ-01-54 revealed a significant main effect of strain [F (1,11) = 20.24, p < 0.001], but no significant effects of drug treatment [F (2,22) = 0.54, p = 0.58] or strain × treatment interaction [F (2,22) = 3.30, p = 0.55]. Consistent with these findings, FMJ-01-54 did not significantly alter total arm entries in either SHR/NCrl or Wistar rats at any administered dose (Fig. 3D).

### FMJ-01-38, but not FMJ-01-54, reduced impulsivity in SHR/NCrl

Consistent with prior findings (de la Peña et al., 2017; de la Peña et al., 2021), vehicle-treated SHR/NCrl showed a reduced percentage of choice for the large, delayed reinforcer, indicating greater impulsive choice (Fig. 4A, C), and an increased frequency of lever presses for the small, immediate reward during the delay period, when such responses were nonreinforced, indicating impulsive action (Fig. 4B, D).

**Figure 4.**
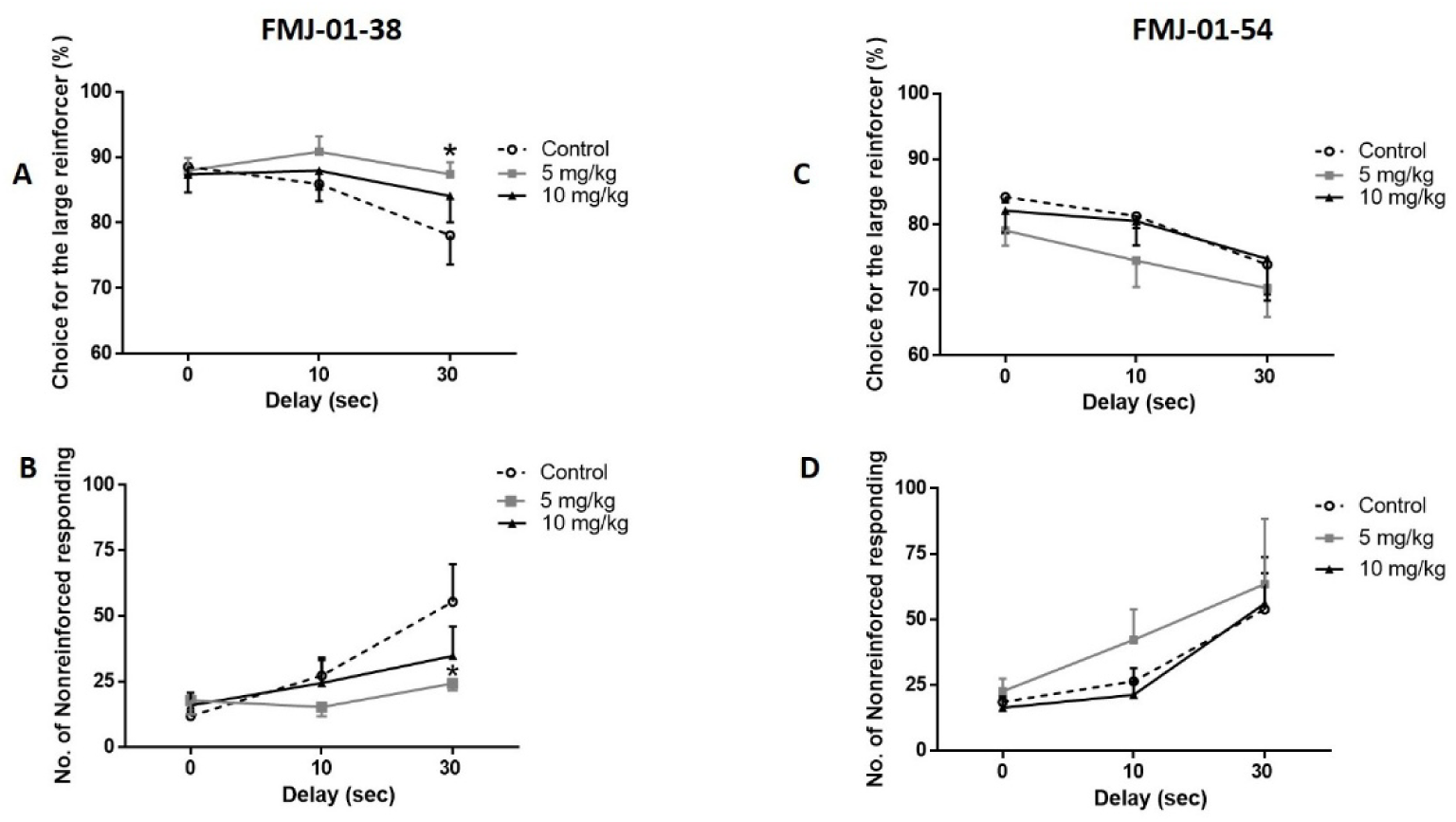
Effects of FMJ-01-38 and FMJ-01-54 on impulsive-like behavior in adolescent SHR/NCrl. Panels A and C show the percentage choice for the larger delayed reward, and Panels B and D show the number of nonreinforced responses for the immediate small reward in the delay-discounting task. Data are expressed as mean ± S.E.M. (n = 10 per group). *p < 0.05 vs. SHR/NCrl control (vehicle-treated) rats.

These effects were most evident when the delay to reward delivery was increased to 30 seconds (Fig. 4A–D). At this 30-second delay, FMJ-01-38 (5 mg/kg, i.p.) significantly improved both impulsive choice and impulsive action in SHR/NCrl (p < 0.05 for both; Fig. 4A–B). In contrast, FMJ-01-54 produced no significant effects on either parameter in SHR/NCrl.

As shown in Fig. 5, vehicle-treated Wistar rats also exhibited impulsive-like behavior, characterized by a decrease in % choice for the larger, delayed reinforcer (Fig. 5A, C) and increased lever pressing during the delay interval (Fig. 5B, D) at the 30-second delay. However, in contrast to the SHR/NCrl results, FMJ-01-38 (5 mg/kg, i.p.) further increased impulsive choice and impulsive action in Wistar rats (p < 0.05; Fig. 5A–B). FMJ-01-54 treatment did not significantly alter impulsive choice or action in Wistar rats.

**Figure 5.**
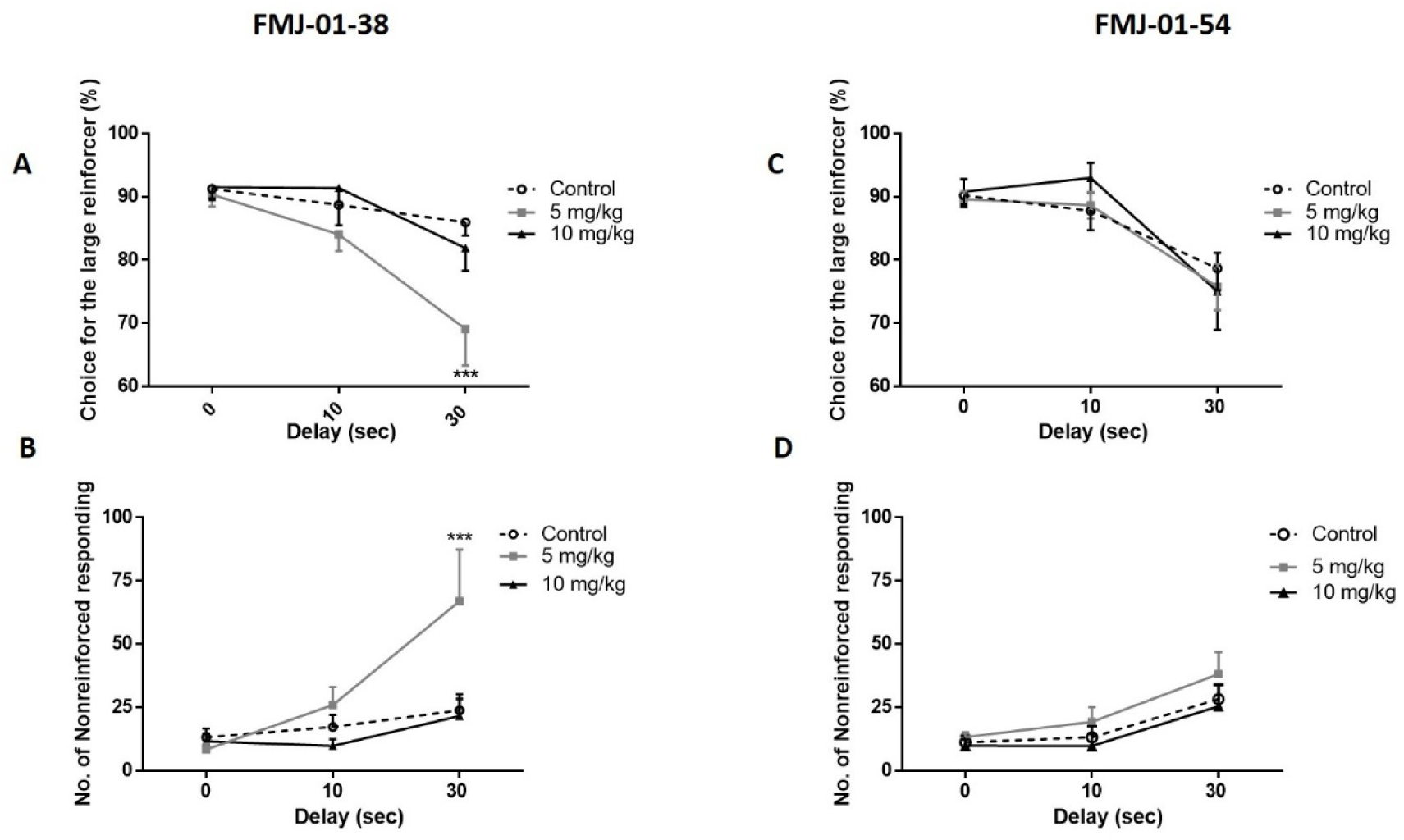
Effects of FMJ-01-38 and FMJ-01-54 on impulsive-like behavior in adolescent Wistar rats. Panels A and C show the percentage choice for the larger delayed reward, and Panels B and D show the number of nonreinforced responses for the immediate small reward in the delay-discounting task. Data are expressed as mean ± S.E.M. (n = 10 per group). ***p < 0.05 vs. Wistar control (vehicle-treated) rats.

## Discussion

The present study examined the effects of two novel D4R ligands, the high-efficacy partial agonist FMJ-01-38 and the antagonist FMJ-01-54, in adolescent SHR/NCrl, a validated model of ADHD, and Wistar rats across four behavioral assays assessing locomotor activity (open field test), attention/recognition memory (novel object preference), attention and working memory (Y-maze test), and impulsivity (delay discounting task). FMJ-01-38 dose-dependently (5–10 mg/kg, i.p.) reduced locomotor hyperactivity and improved spontaneous alternation behavior in SHR/NCrl; at 5 mg/kg, it further enhanced novel-object preference and reduced impulsive choice and action in the delay-discounting task, indicating amelioration of ADHD-like behaviors and improved cognitive performance. In contrast, FMJ-01-54 produced dose-dependent improvements in Y-maze and novel-object preference performance, but did not reduce locomotor hyperactivity or impulsivity in SHR/NCrl, suggesting a selective cognitive-enhancing effect. FMJ-01-38 enhanced novel object preference only at the 5 mg/kg dose in Wistar rats, whereas FMJ-01-54 produced no significant behavioral effects.

Previous studies have demonstrated the cognitive-enhancing effects of partial D4R activation as well as potential mechanisms underlying attention- and impulse control-improving actions of the D4R partial agonist, RO-10-5824 (Nakazawa et al., 2015; Powell et al., 2003). The present findings extend these observations by showing that FMJ-01-38 not only enhances cognition in unimpaired animals but, importantly, attenuates hyperactivity, inattention, and impulsivity in the SHR/NCrl ADHD animal model. As previously noted, SHR rats exhibit markedly reduced *Drd4* gene expression and D4R protein synthesis in the PFC (Li et al., 2007) accompanied by a hypoactive dopaminergic system characterized by reduced vesicular dopamine storage and release (Russell, 2002; Sagvolden et al., 2005). Consistent with previous reports that full D4R agonists (e.g., ABT-724, A-412997) alleviated SHR hyperactivity and cognitive deficits (Browman et al., 2005; Yin et al., 2014), the behavioral improvements observed with FMJ-01-38 likely reflect restoration of PFC inhibitory regulation via submaximal D4R activation and compensation for reduced dopaminergic tone, thereby stabilizing cortico-striatal signaling known to be dysregulated in ADHD (Durston et al., 2011; A. Nikolaidis et al., 2022; Volkow et al., 2009). In SHR rats, where PFC D4R density is reduced, FMJ-01-38 likely acts predominantly as an agonist, engaging Gi/o-coupled signaling and modulating downstream cAMP–PKA and ERK–MAPK pathways implicated in attention and executive control (Funk et al., 2012; González et al., 2012; Peng et al., 2010). Through these mechanisms, FMJ-01-38 may restore excitatory–inhibitory balance in the PFC, normalize cortical output, and reduce behavioral manifestations of hyperactivity, inattention, and impulsivity in SHR/NCrl. The precise cellular and molecular mechanisms mediating these effects warrant further experimental investigation. In contrast, in Wistar rats with presumably intact dopaminergic tone, FMJ-01-38 would typically be expected to act as a functional antagonist, producing little or no behavioral effects consistent with the state-dependent nature of partial agonists (Nakazawa et al., 2015; Powell et al., 2003). Interestingly, FMJ-01-38 further increased impulsivity in Wistar rats, suggesting that its antagonistic effect may have subtly disrupted PFC dopaminergic balance, thereby impairing inhibitory control.

Nonetheless, FMJ-01-38 improved novel object preference in Wistar rats suggesting that partial D4R activation may also improve cognition beyond ADHD-related pathology. Although the compound reduced total arm entries in the Y-maze test, the lack of corresponding changes in open-field locomotion indicates that this effect more plausibly reflects attenuated exploratory behavior, rather than drug-induced sedation or locomotor suppression. Taken together, these results suggest that partial D4R modulation in Wistar rats can differentially influence inhibitory control and motivational processes while simultaneously preserving, or in some cases enhancing, D4R-dependent cognitive functions.

Aside from partial agonists, the therapeutic potential of D4R blockers has also been reported. Selective D4R antagonists such as L-745,870 and U-101,958 reduced hyperactivity in dopamine-depleted or lesion-based ADHD models without affecting locomotor activity in control animals (Zhang et al., 2002). In this study, however, FMJ-01-54 did not attenuate SHR/NCrl’s hyperactivity, likely reflecting model-specific or drug-related differences. Notably, 6-OHDA lesions broadly affect dopaminergic projections and are not restricted to the D4R subtype (Browman et al., 2005). Moreover, previously used D4R antagonists exhibit off-target serotonergic and sigma receptor activity (Lee et al., 2008; Zhang et al., 2002). Nonetheless, FMJ-01-54 improved attention and recognition memory in SHR/NCrl, representing, to our knowledge, the first evidence of a D4R antagonist enhancing cognition in a validated ADHD model. Notably, L-745,870 has been shown to improve working memory in other studies, with an inverted U-shaped dose–response relationship, enhancing performance only in subjects with lower baseline cognitive function (Zhang et al., 2004). The selective effects of FMJ-01-54 in SHR/NCrl, but not in Wistar rats, are consistent with this observation. Furthermore, FMJ-01-54 did not affect locomotion or impulsivity, suggesting that D4R antagonism exerts selective cognitive-enhancing effects that emerge primarily under conditions of dopaminergic dysfunction. Further studies are needed to elucidate the molecular and neurophysiological mechanisms underlying the effects of FMJ-01-54 on ADHD-related cognitive and attentional/working memory deficits.

From a translational perspective, these findings suggest the potential of D4R ligands as novel, non-stimulant therapeutics for ADHD. Notably, FMJ-01-38 improved ADHD-like behavioral deficits in SHR/NCrl while producing minimal effects in control animals, consistent with the concept of pathology-selective therapy, i.e., treatments that normalize dysregulated neural function without altering normal brain activity (Arnsten, 2006; Vaidya et al., 1998). Unlike psychostimulants that broadly increase catecholamine levels (de la Peña et al., 2015), D4R partial agonists such as FMJ-01-38 may provide targeted modulation of fronto-striatal circuits, thereby reducing abuse liability and peripheral adverse effects. Moreover, by producing submaximal receptor activation, D4R partial agonists such as FMJ-01-38 may confer therapeutic benefits while reducing the behavioral and physiological effects typically observed with full D4R stimulation. In parallel, the selective cognitive benefits of FMJ-01-54 suggest that D4R antagonism could complement partial agonism by improving attentional or working memory function through distinct mechanisms. Collectively, these results strengthen the therapeutic relevance of D4R modulation and warrant further investigation into receptor-state dynamics, signaling mechanisms, and long-term adaptations underlying D4R-targeted pharmacologic interventions.

The present study has a few limitations that warrant consideration. First, the behavioral effects of D4R drugs were assessed at only two doses, necessitating broader dose–response studies to better define their therapeutic and adverse profiles. Second, as ADHD is a chronic disorder requiring long-term treatment, future work should evaluate the sustained efficacy and safety of these compounds under chronic dosing regimens to identify potential drug-induced neuroadaptive changes. Third, the study was limited to male adolescent rats, precluding assessment of sex- or age-dependent differences in D4R function. Fourth, although SHR/NCrl are validated ADHD models, they cannot fully capture the complexity of human ADHD symptoms such as inattention and impulsivity. Finally, the study relied primarily on behavioral parameters without direct neurochemical or molecular correlates; thus, the mechanistic basis of the observed effects, such as modulation of PFC D4R signaling, dopamine release, or downstream signaling cascades, remains to be elucidated.

In conclusion, this study demonstrated that the D4R-targeting compounds FMJ-01-38 and FMJ-01-54 produced distinct yet significant behavioral effects in the SHR/NCrl ADHD animal model. These findings support the therapeutic potential of D4R modulation, suggesting that both partial agonism and antagonism may confer symptom-selective benefits in ADHD through distinct mechanisms. Overall, the results suggest the D4R as a promising molecular target for developing safer and more specific non-stimulant treatments for ADHD.

## Acknowledgments

This research was supported by the National Institutes of Health/National Institute on Drug Abuse (NIH/NIDA) under Grant DP1DA058385 and in part by the U.S. Health Resources and Services Administration (HRSA) under Grant D34HP45723, and the Loma Linda University School of Pharmacy.

## Author contributions

All authors contributed to the study conception, design and execution. ICD, TMK, and CAB conceptualized the study design. Material preparation was performed by MAA, TL, TMK and CAB. Data collection and analysis were performed by ICD, SA, and AA. The first draft of the manuscript was written by ICD and all authors provided feedback on earlier versions of the manuscript. All authors read and approved the final manuscript.

## Data availability

Data are made available upon reasonable request

## Declarations

### Ethical approval

All procedures were conducted in accordance with the Public Health Service Policy on Humane Care and Use of Laboratory Animals and the Guide for the Care and Use of Laboratory Animals and were approved by the Loma Linda University Institutional Animal Care and Use Committee (IACUC #24-015).

## Consent to participate

Not applicable.

## Consent to publish

Not applicable.

